# Long non-coding RNA UCA1 affects chromatin remodeling via SMARCA2-containing SWI/SNF complex in human colorectal cancer

**DOI:** 10.1101/2024.09.27.615411

**Authors:** Bernadette Neve, Elsa Hadj Bachir, Belinda Duchêne, Mouloud Souidi, Martin Figeac, Jean-Pascal Meneboo, Emmanuelle Leteurtre, OrgaRES consortium, Nicolas Jonckheere, Audrey Vincent, Isabelle Van Seuningen

## Abstract

Differential lncRNA expression has been correlated to clinical characteristics of colorectal cancers (CRC), which are the second leading cause of cancer-related deaths. LncRNA UCA1 plays a role in epigenetic gene regulation in diverse cancers. We studied CRC cell properties in CRISPR/Cas9 HT29-derived models and, interestingly, UCA1-depleted HT29 cells presented an increased stem-cell phenotype. We show that loss of UCA1 affected SWI/SNF chromatin remodeling complexes, which are previously shown to be involved in maintaining stem-cell properties. Not only was UCA1 permissive for induced SWI/SNF subunit SMARCA2 (BRM) expression upon chemo-drug treatment, but it also affected subunit compositions of SWI/SNF complexes by direct interaction of UCA1 with both ATP helicase BRM and BRG1. UCA1 is known to stimulate proliferation and decrease apoptosis, and we here show that it can restrain cells from a stem cell phenotype. The dual action of UCA1 revealed in this study highlights the complex actions of lncRNAs in cancer.

## INTRODUCTION

Colorectal cancers (CRCs) have become the second leading cause of cancer-related deaths in Europe.^1^ Acquired chemoresistance and the development of metastatic lesions, associated with the presence of cancer stem-cells, are a cause of poor survival rates. On average the 5-year survival rate is 14% for patients with metastatic CRC.^2^ Besides expression of protein-coding genes, differential expression of diverse long non-coding RNA (lncRNA) transcripts has also been correlated with different clinical CRC characteristics and with molecular phenotypes.^3^

UCA1 is a lncRNA that is up to 2.7 kb in length and that is implicated in diverse cancers including colorectal cancer.^4–8^ UCA1 resides both in the cytoplasm and nucleus of cancer cells. Recently, it was shown that N6-adenosine methylation of UCA1 increases its stability via interaction with IGF2BP2 in the cytoplasm.^9^ Several studies suggested that cytosolic UCA1 regulates miRNA-mediated decay of genes by acting as a competing endogenous RNA.^4^ For example, the short isoform of UCA1 (1.4 kb) can inhibit apoptosis by binding miR-27a-5p and regulating UBE2N expression in ovarian cancer cells.^10^ That study also reported that a long isoform of UCA1 is mainly present in the nucleus. Nuclear UCA1 stimulates, and is crucial for, cell cycle progression.^5,6^ Indeed, UCA1 is known to negatively regulate transcription of cell cycle inhibitors p21cip and p27Kip1, ^11–13^ and to stimulate transcription of cyclin D1 at the promoter level.^14^ Epigenetic regulation of transcription by UCA1 is also highlighted by its interaction with enhancer of zeste homolog 2 (EZH2) regulating histone 3 tri-methylation (H3K27me3) ^11–13^ and by the interaction with chromatin insulator CCCTC-binding factor (CTCF).^15^ Interestingly, the presence of UCA1 resulted in less binding of Brahma related gene 1 (BRG1/SMARCA4) at p21cip promoter locus in bladder cancer cells.^16^ In fact, RNA-pull down in these cells showed interaction of UCA1 with BRG1-containing SWI/SNF complexes. In our study, we investigated gene regulation by UCA1 in colorectal cancer cells.

Several studies have shown a role for SWI/SNF chromatin remodeling complexes in cancer.^17–19^ Different SWI/SNF remodeling complexes each harbor exclusively either ATP-dependent helicase BRG1 (*SMARCA4*) or Brahma (BRM, *SMARCA2*). The assembly with other sub-units is dynamic during cellular differentiation and distinguishes different complex classes, such as PBAF (Poly-Bromo-Associated Factors; exclusively with BRG1), canonical and non-canonical BAF complexes (BAF; BRG1- or BRM-associated Factors).^20,21^ The PBAF complex containing BRG1, core subunits, and the specific subunits polybromo 1 (PBRM1), AT-rich interaction domain (ARID)2 and Bromodomain containing (BRD) 7, is implicated in the development and maintaining of cellular identity.^22–24^ Down regulation of human PBRM1 from the PBAF complex in prostate cancer cells resulted in more sphere formation, which signifies the presence of more stem cells.^23^ Interestingly, cells that are drivers of carcinogenesis are associated with multipotent stem-cell phenotypes.^25^ In general, the SWI/SNF complex that is frequently associated to stemness (esBAF), is a BRG1-containing complex assembled with core subunits in combination with supplementary subunits SMARCE1, SMARCC2 and ARID1A.^26,27^ Other BAF-complexes are tissue and cell type specific, for example in progenitor cells they specifically harbor SMARCD3 and SMARCD1 subunits (cardiac and neuronal progenitors, respectively). ^28,29^ The presence of either BRG1 or BRM does not classify the different SWI/SNF complexes, although BRG1 and BRM may associate with ‘actively marked’ and ‘repressively marked’ chromatin regions, respectively.^30^ Mutations of SWI/SNF subunits genes are frequently associated with cancer^31–33^ and may evoke druggable vulnerabilities ^34,35^ underlining the relevance of our study on this chromatin remodeling family.

A relation of lncRNA UCA1 with SWI/SNF-complexes is previously shown in breast cancer cells where UCA1 expression is inhibited by BAF-subunit ARID1A,^36^ and also in bladder cancer cells where an interaction of UCA1 with BRG1-containing SWI/SNF-complexes is shown.^16^ In this study, we showed that UCA1 depletion was associated with a stem cell phenotype. Transcript expression of several SWI/SNF subunits, in particularly SMARCA2, was affected by UCA1 that was increased by cancer therapy drugs 5-Fluorouracil and Oxaliplatin treatment in our colorectal HT29 cell model. Moreover, we showed a direct interaction of UCA1 with both BRG1 and BRM, suggesting a role of UCA1 in the balance of the different SWI/SNF complexes. Indeed, we revealed a role of UCA1 in the shift of BRG1-containing SWI/SNF complex from PBAF to canonical BAF.

## RESULTS

### LncRNA UCA1 expression is increased in CRC tumor tissue

High UCA1 expression is correlated with bad disease prognosis in diverse colorectal cancer studies.^4–8^ We measured UCA1 expression in normal tumor-adjacent (NAT) compared to tumor tissues from 13 patients of the Lille university hospital (OrgaRES study) and showed that UCA1 expression was indeed significantly higher in tumor samples (Figure 1A, *p* <0.01).

**Figure 1.**
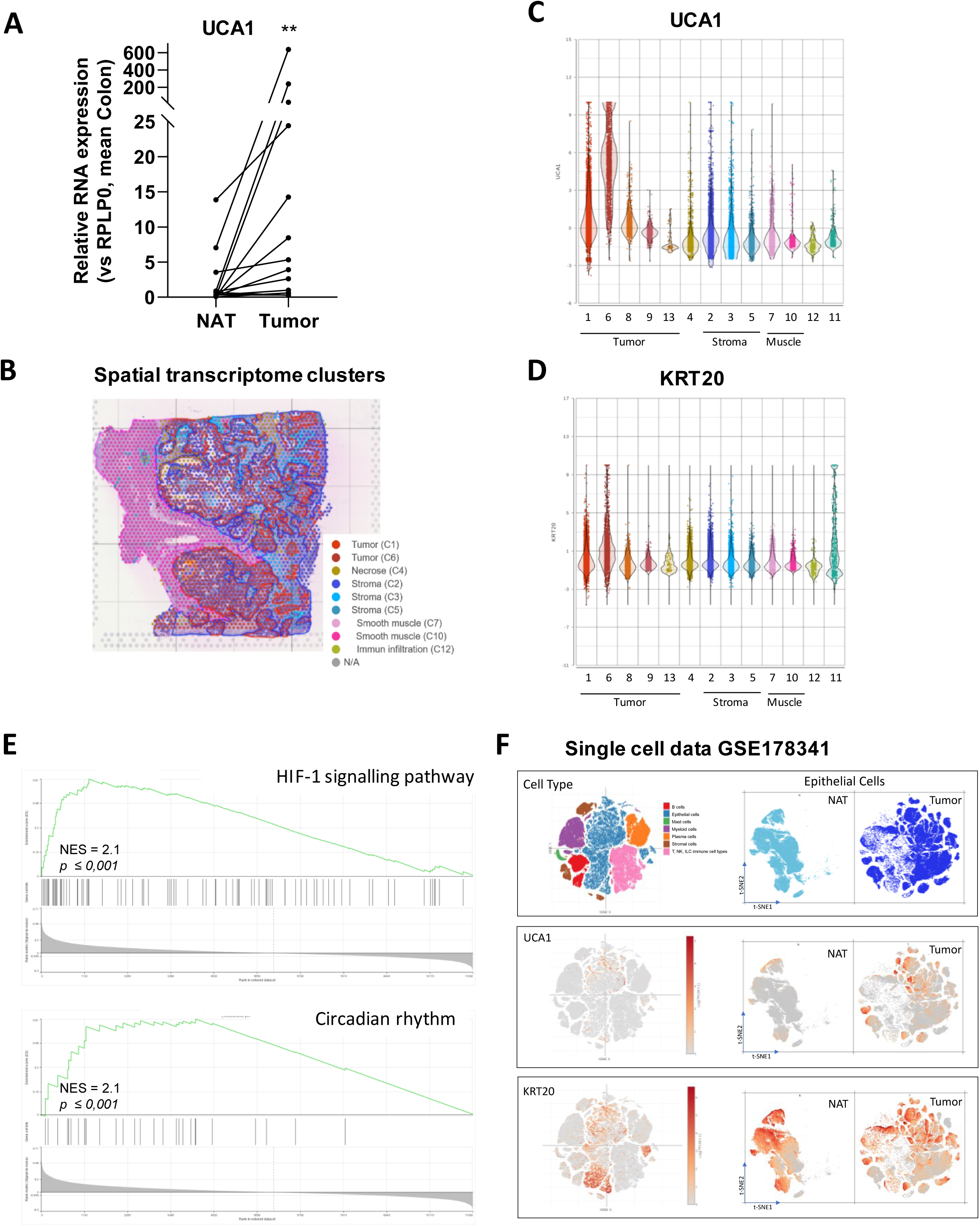
LncRNA UCA1 expression is increased in CRC tumor tissue. (A) UCA1 expression in cryo-preserved CRC tumors and normal tumor-adjacent colon tissues (NAT) from 13 patients (**p <0.01 of paired t-tests on ratios). (B) Visium spatial transcriptome analysis of 12 CRC tumor tissues with an example of Hematoxylin-Eosin staining with pathologist-assigned tumor regions encircled in red and Visium capturing spots colored based on the 13 graph-based expression clusters identified within all tissue samples (dots legend). Cluster 1, 6, 8, 9 and 13 were assigned as tumor tissue by our pathologist (dots colored Red to Orange). (C and D) Spatial distribution of Log2 expression of UCA1 and KRT20 transcripts observed in the graph-based clusters. (E) Enrichment profiles of cluster 6 genes compared to cluster 1, 8, 9 and 13 for the HIF-1 signaling pathway and circadian rhythm (both NES =2.1, *p*-value ≤0.001). (F) t-SNE projections of single colorectal cells colored by cell-type, and of epithelial cells split on the basis of disease ontology (normal tumor-adjacent (NAT) and tumor) (top) both colored for log2-expression of UCA1 and KRT20, respectively.

In addition, we studied how UCA1 expression was distributed in heterogenous tumor tissues by spatial transcriptome analysis of 12 CRC tumor samples. Our pathologist annotated the tumorous tissue regions on the HE-colored slide. Combined principal component analysis of the samples in Partek resulted in assignment of 13 graph-based expression clusters (Figure 1B). UCA1 expression was enriched and identified as top biomarker of tumor cluster 6 (Figure 1C; 29.6-fold increased expression, FDR ≤0.001). This cluster was also enriched for enterocyte marker KRT20 (Figure 1D; 11.0-fold increased expression, FDR ≤0.001).

Compared to other clusters that were annotated as tumor enterocyte tissue by our pathologist (Cluster 1, 8, 9, 13), cluster 6 was enriched for HIF-1 signaling and circadian rhythm pathways ((Figure 1E, for both GSEA normalized enrichment score (NES) =2.1, *p*-value ≤0.001). Within cluster 6, UCA1 significantly correlated with 502 genes, including CCND2 and CDK1 (absolute R ≥0.20, FDR ≤0.001). In line with previous implication of UCA1 with cell cycle progression, these correlated genes showed a pathway enrichment for cell cycle and DNA replication (respectively, NES = 12.3 and 9.5, *p*-value ≤0.001).

To analyze UCA1 expression at a single cell level we studied publicly available single cell expression data (Figure 1F). Similar to our spatial transcriptome results, we observed that UCA1 was higher expressed in tumor epithelial cells with 1.2-fold higher expressed in the high KRT20-expressing CRC epithelial cells (23063 cells with KRT20 log2 expression >2 compared to 90530 cells log2 ≤2, *p* <0.001). These results indicate UCA1 expression was increased in a subclass of tumor cells.

### Loss of UCA1 expression induces a stem cell-like phenotype

To study effects of stable UCA1 depletion on CRC cell properties, we derived HT29 cell clones with a genomic deletion at exon 1 or exon 2 of the *UCA1* gene locus using CRISPR/Cas9. We selected HT29iCAS9-delta UCA1 clones showing an alteration at the UCA1 genomic region and a loss of UCA1 expression in qPCR analysis (Figure S1A). Similar to a lot of lncRNA expressions, the HT29-derived control (ICP) cells expressed very low UCA1 levels (Figure S1B), however, treatment of the cells with the chemo-drug 5-Fluorouracil (5-FU) increased UCA1 levels. We studied alternative UCA1 transcript expression by RNA sequencing (Figure S1B). In the control ICP cells, UCA1 variants 213 (size 2.7kb), 216 (2.6kb), 214 (1.6kb), 243 (1.8kb) and 206 (1kb) significantly increased upon 5-FU treatment (Figure S1B, FDR <0.01). In both the dUCA1E1 and dUCA1E2 cells, UCA1 was neglectable and only variant 207 lacking exon 2 (1.2kb) was significantly induced by 5-FU in the dUCA1E2 cells (FDR <0.001). When we assessed previously reported UCA1 target genes,^4^ we observed that SIRT1, SF1, GLS2, PFKFB2, HK2, BCL2, PBX3 and ZEB1 had a suggestive difference of 5-FU-induced expression changes in dUCA1 cells (Figure S1C).

Next, we performed tumorosphere formation assays to analyze stem cell properties in our model. With ELDA assays, we showed that dUCA1 cells had a significantly increased proportion of stem cells compared to the control ICP cells (Figure 2A; frequency of 0.18 for ICP, 0.29 for dUCA1E1 (vs ICP **p<0.01) and 0.26 for dUCA1E2 (vs ICP *p<0.05)). When multiple cells were seeded the formed spheres tended to merge (Figure 2B). Analysis of individual cell culture assays not only confirmed that the stemness capacity was increased, but also showed that spheres from both dUCA1E1 and dUCA1E2 cells were increased in size compared to ICP cells (Figure 2C; 120 and 122% of ICP cells (surface area 3958±684 µm2, *p<0.05), and Figure 2C; representative PGC-images of measured spheres). These results suggest a novel role for UCA1 in cellular identity.

**Figure 2.**
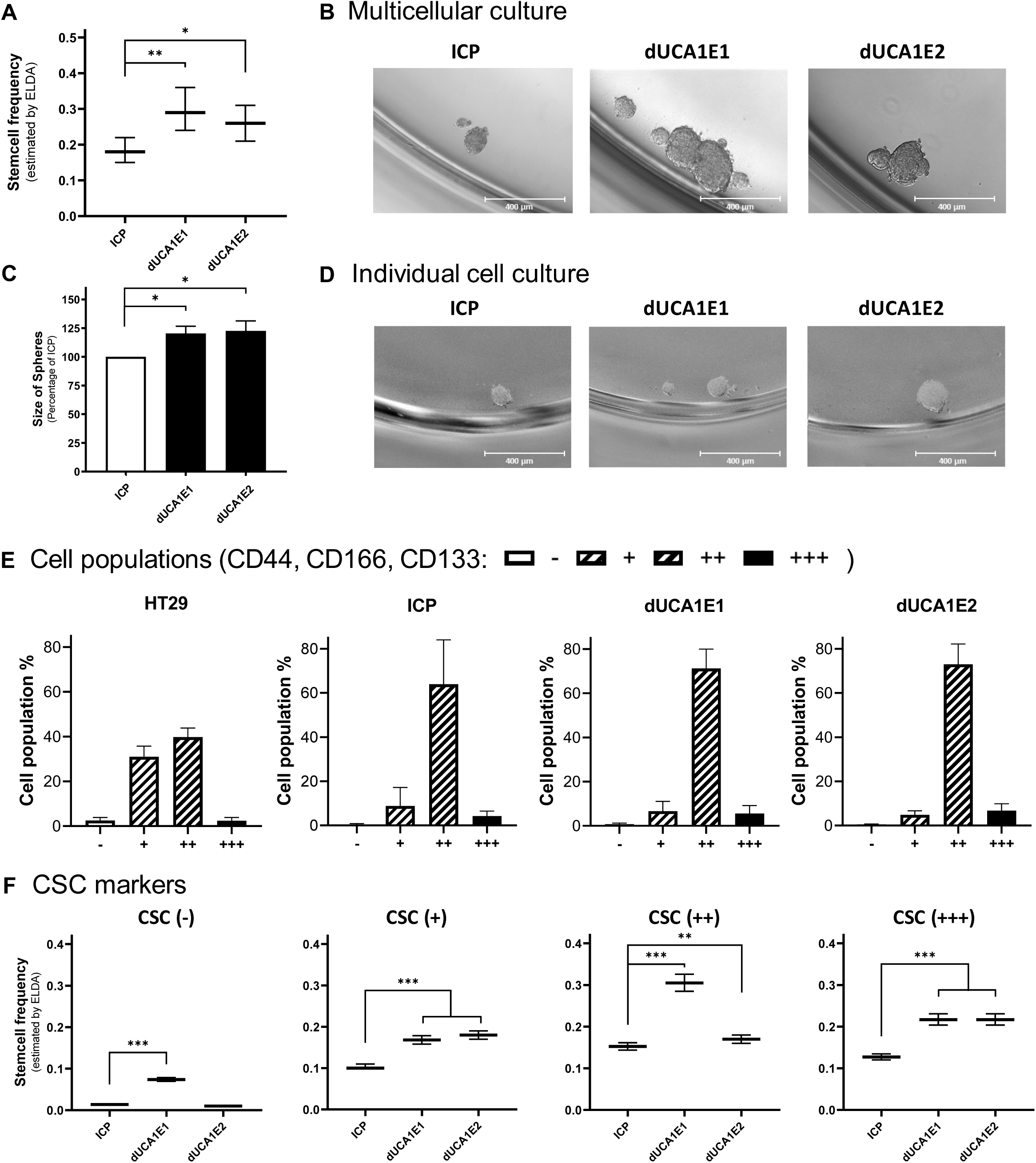
Loss of UCA1 expression induces a stem cell-like phenotype. (A) Ability of HT29-derived cellular clones (control ICP, dUCA1E1 and dUCA1E2) to form spheres. ELDA estimated stem cell frequencies (n=5, *p <0.05, **p <0.01). (B) Representative images of non-adherent multicellular culture for 7 days (Microscope Leica DMi8). (C) Size of spheres measured after individual culture of ICP, dUCA1E1 and dUCA1E2 cells in non-adherent conditions (n=7, *p <0.05). (D) Representative Phase Gradient Contrast (PGC) images of non-adherent individual sphere culture for 11 days (Microscope Zeiss Cell discoverer 7) (E) Percentage of different cell populations in HT29, control ICP, dUCA1E1 and dUCA1E2 cells (n=3-9). Cells were labelled for expression of stem cell markers CD44, CD133 and CD166 and sorted by FACS into a pool of cells expressing no markers (−), one of three (+), two of three (++), or all markers (+++). (F) Different pools of cells expressing CSC markers were analyzed for their ability to form spheres by ELDA (n=3, **p <0.01, ***p <0.001).

### The ICP and dUCA1 cells have similar stem cell markers expression

The ability to form spheres is correlated with the expression of stem cell markers CD44, CD133 and CD166.^37^ Therefore, we studied by flow cytometry the distribution of cells expressing these stem cell markers. Since control ICP cells had a low population of stem cell marker-negative cells, we also included the original HT29 cells in our analysis. The control ICP cells have a lower stem cell markers-negative (−) or simple positive population (+) and a higher proportion of cells with two or three stem cell markers (++/+++) compared to original HT29 cells (Figure 2E; (−) 2.5% vs 0.4%, (+) 31% vs 9%**, (++) 40% vs 64%** and (+++) 2.3% vs 4.2% for the different populations of HT29 vs ICP cells, respectively, \*\**p* <0.01). However, no difference in cell population distribution was observed between ICP and dUCA1 cells.

We analyzed the capacity to form spheres of cells with no markers (−), one of three (+), two of three (++), or all three markers (+++) expression after FACS-sorting. Indeed, ICP cells expressing one or more stem cell markers presented a higher ELDA estimated stem cell frequency based upon tumor-sphere formation (Figure 2F). Stem cell frequency varied in the dUCA1 cells, but both dUCA1E1 and dUCA1E2 cells with one or more markers at the surface had a significantly increased stem cell frequency compared to the same population of ICP cells. From these results we concluded that UCA1 depletion evoked intrinsic phenotype changes independent of CSC marker expression.

### UCA1 depletion affects the SWI/SNF subunit SMARCA2 expression

We further analyzed our dUCA1 model for differentially expressed genes that could be related to the differences in stem cell properties. We observed 158 genes that showed a significantly different change in expression upon 5-FU treatment in dUCA1 cells compared to ICP cells (Figure 3A, FDR <0.05). No obvious stem cell candidates were revealed, but we observed a suggestive lack of increased expression upon 5-FU (ICP vs dUCA1) for several subunits of the SWI/SNF chromatin remodeling complex (Figure 3B).

**Figure 3.**
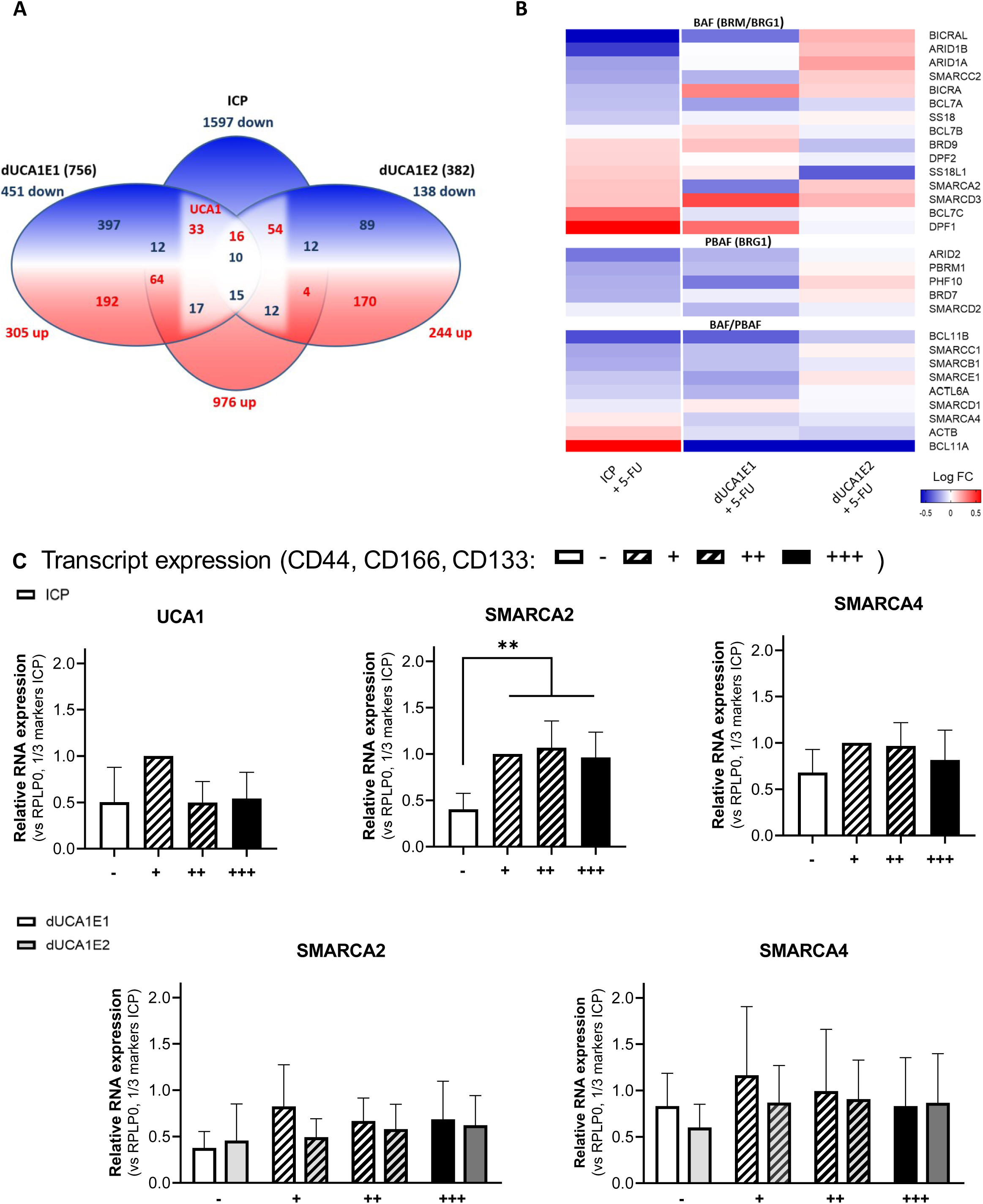
Decreased SMARCA2 expression correlates with UCA1 depletion. (A) RNAseq analysis of control ICP, dUCA1E1 and dUCA1E2 cells treated for 72h with or without 2 µM 5-Fluorouracil (5-FU). Numbers represent differentially expressed genes in ICP cells upon 5-FU treatment compared to dUCA1 cells with 5-FU (FDR<0.05). White area represents opposite gene regulation in dUCA1 cells (158 genes). (B) Heatmap of SWI/SNF-subunit gene expressions in ICP cells upon 5-FU treatment compared to dUCA1 cells with 5-FU. (C) Relative transcript expression of UCA1, SMARCA2 and SMARCA4 in different cell populations (see legend Figure 2E). Since (−) cell populations were small, expressions were calculated relative to the (+) cells (n=5, \*\**p* ≤0.01).

The presence of different SMARCA2 (BRM)- and SMARCA4 (BRG1)-containing complexes has been associated with cancer stem-cell phenotypes, therefore we assessed UCA1, SMARCA2 and SMARCA4 expression in stem cell markers sorted cells (Figure 2C). For UCA1, no different expression was observed in the different ICP cell populations. For SMARCA2, we observed in ICP cells an increased transcript expression in populations having one or more stem cell markers compared to stem cell markers negative cells (Figure 2C; *p* ≤ 0.01). SMARCA4 expression did not vary between the different ICP cell populations. Interestingly, dUCA1E1 and E2 cells were different compared to ICP cells as no changes of SMARCA2 transcript levels in the stem cell markers cell populations were observed. These results indicate that UCA1 interfered with SMARCA2 transcript levels.

### The signature of SWI/SNF subunit expressions defines a CRC risk group

To assess the importance of SWI/SNF subunits on survival of CRC patients, we assessed their expression in the TCGA COAD-READ study. Based on subunits expression levels, multivariate analysis using SurvExpress defined two distinct risk groups for overall survival (Figure 4C; HR =2.43, Confidence interval: 1,58-3,74, *p* <0.0001). The generated box-plot showed lower expression of BAF-related subunits (first 16 listed) and higher expression of PBAF-related subunits (number 17 - 21) in the low-risk group compared to the high-risk group (significantly different expression between groups ranging from *p* <0.02 for SMARCA2 to *p* <2.0e-10 for BRD9). In addition, multivariate analysis of TCGA data on other digestive tract cancers showed that the expression signature of SWI/SNF subunits also defined significantly different risk groups in pancreatic adenocarcinoma (PAAD, HR =2.6, *p* <0.0001), esophageal carcinoma (ESCA, HR =3.5, *p* <0.0001), stomach adenocarcinoma (STAD, HR =1.87, *p* <0.001) and in the stomach and esophageal carcinoma studies (STES HR =1.82, *p* <0.001).

**Figure 4.**
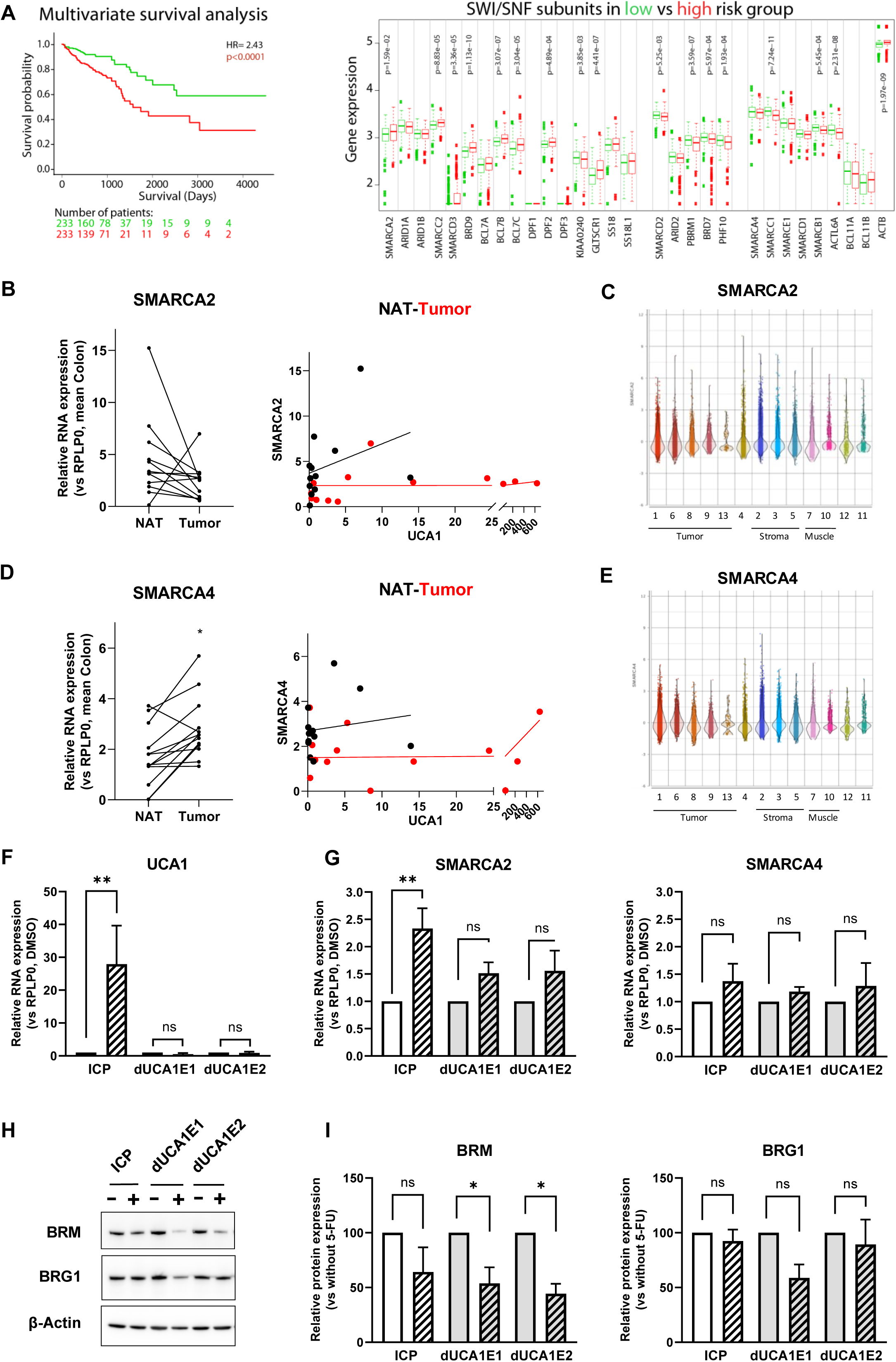
UCA1 stabilizes SMARCA2 protein expression (BRM) after 5-FU treatment. (A) Kaplan-Meier survival plot of SWI/SNF sub-units expression in COAD-READ, with risk groups defined by median Cox prognostic indexes, showing that the expression signature is correlated with overall survival (HR=2.43; CI 1,58-3,74; *p*<0.0001). Box plot depicting gene expression of SWI/SNF subunits in the low (green) and high (red) risk group. Indicated above the boxes are significant *p*-values of differential expression between risk groups (B) SMARCA2 transcript expression in tumor and NAT tissues. Correlation between SMARCA2 and UCA1 expression analyzed by simple linear regression. (C) SMARCA2 transcript distribution (Log2-expression) in graph-based clusters of Visium spatial transcriptome analysis. (D) SMARCA4 transcript expression in tumor and NAT tissues. Correlation between SMARCA4 and UCA1 expression analyzed by simple linear regression. (E) SMARCA4 transcript distribution (Log2-expression) in graph-based clusters of Visium spatial transcriptome analysis. (F and G) UCA1, SMARCA2 and SMARCA4 transcript expression in control ICP, dUCA1E1 and dUCA1E2 cells treated for 72h with 2 µM 5-FU (striped bars; n=5, **p<0.01). (H) BRM and BRG1 protein expression (encoded by *SMARCA2* and *SMARCA4*, respectively) in control ICP, dUCA1E1 and dUCA1E2 cells treated for 72h with 2 µM 5-FU. (I) Relative expression of BRM and BRG1 protein compared to ß-actin on the western blots, quantified by ImageJ analysis (n=3, * p< 0.05).

To further analyze the association of UCA1 expression with SWI/SNF complexes, we evaluated the correlation with SMARCA2 and SMARCA4 in the COAD-READ dataset. UCA1 expression was significantly negatively correlated with SMARCA2 expression (Figure S2A; R =-0.26 and *p* <0.0001), but not correlated with SMARCA4. Interestingly, similar correlations of UCA1 expression with SMARCA2 expression was observed in PAAD, ESCA, STAD and STES study (Figure S2B, S2C, S2D and S2E, respectively).

We also assessed expression of the two ATP-dependent helicases of SWI/SNF complexes, SMARCA2 and SMARCA4, and the correlation with UCA1 expression in our OrgaRES tissue samples. Here, transcript expression of SMARCA2 was not significantly different in the tumor compared to NAT samples and there was no evidence for a correlation with UCA1 expression (Figure 4B). In our spatial transcriptome analysis, SMARCA2 showed a general distribution between the different graph-based expression clusters (Figure 4C). SMARCA4 expression was significantly increased in tumor compared to NAT tissue (Figure 4D, *p* < 0.05), but no correlation with UCA1 expression was observed in these samples, nor in our spatial transcriptome analysis (Figure 4D and 4E). The fact that UCA1 was mainly expressed in cells with high expression of the enterocyte marker KRT20, whereas SMARCA2 had a broader expression suggest the observed correlation between UCA1 and SMARCA2 in bulk analysis was based on the presence of different cancerous tissue types.

### Loss of UCA1 in HT29-derived cells modulates SMARCA2 expression

To further study the relation of UCA1 with SMARCA2 and SMARCA4, we performed qPCR and Western blot analysis in our dUCA1 cell line model. In the control ICP cells, UCA1 had a low expression level under standard culture conditions, which was significantly increased upon treatment with 2 µM 5-FU for 72h (Figure S1 and 4F). In both the dUCA1E1 and dUCA1E2 cells, UCA1 can no longer be detected by qPCR. Upon treatment with 5-FU, the expression of SMARCA2 transcript was significantly increased in the UCA1 expressing control ICP cells, but not in the dUCA1 cells (Figure 4G). The expression of SMARCA4 transcript was not significantly affected by the 5-FU treatment, nor by the loss of UCA1 expression. While the SMARCA2 transcript expression increased upon 5-FU treatment, the SMARCA2 protein level (BRM) remained similar in 5-FU-treated cells compared to control cells (Figure 4H and 4I). This may result from an initial degradation of BRM after 5-FU treatment that was not entirely compensated by increased transcript translation. Indeed, in the absence of UCA1 there was no change of SMARCA2 transcript level and a decreased BRM protein level was observed in dUCA1E1/E2 cells, suggesting protein degradation (Figure 4I). Analyzing SMARCA4 (BRG1) protein level expression under these conditions showed no changes under the different conditions (Figure 4H and 4I). Similar trends were observed upon treatment of the cells with 2 µM of the chemo-drug oxaliplatin (Figure S3). These results confirm our initial findings with sorted stem cells, which indicated that UCA1 interferes with SMARCA2 at a transcription level.

### UCA1 physically interacts with the ATP-dependent helicases BRM and BRG1

Loss of UCA1 evoked less SMARCA2 (expression and protein) and more stem-like cells, and in contrast, cells bearing stem cell markers showed higher SMARCA2 expression. SWI/SNF Complexes that are associated with a stem cell phenotype harbor mainly the other helicase BRG1 (SMARCA4). We hypothesized that UCA1 may also physically affect the subunit balance of SWI/SNF-complexes. Therefore, we first analyzed direct physical interaction of UCA1 with BRM and BRG1 proteins in the colorectal HT29 cells using a Duolink proximity ligation assay adapted for ncRNA-protein interactions. In these cells both BRM- and BRG1 interacted with UCA1 (Figure 5A-5D). Moreover, increasing UCA1 expression by treating the cells with 2 µM 5-FU for 24h resulted in an increased number of interactions, with, in particularly, more nuclei having multiple spots (Figure 5A-5D). Some interactions were also observed in the cytoplasm. In addition, our RNA-IP analysis showed interaction of UCA1 with BRM- and BRG1-complexes (Figure 5E), also suggesting that UCA1 interfered with these complexes on a protein level.

**Figure 5.**
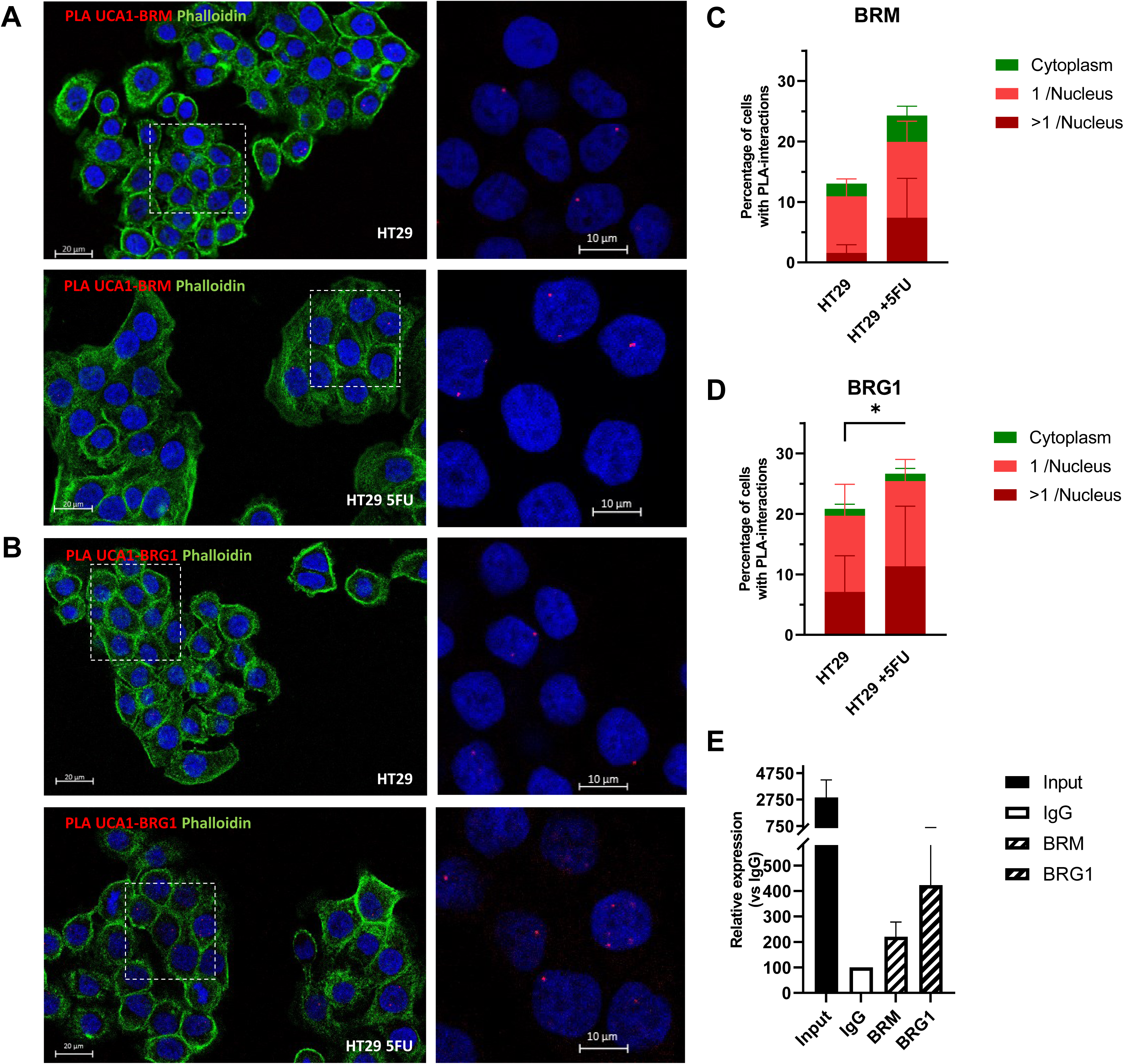
UCA1 physically interacts with BRM and BRG1 in HT29 cells. (A and B) Representative confocal microscope images of PLA assays of HT29 cells with or without 2 µM 5-FU for 24h showing nuclear staining (Dapi, blue), actin (Phalloduin, green) and dotted interaction signals (PLA, red) of UCA1 with BRM (A) and with BRG1(B). Enlarged sections are marked by dotted white lines. (C and D) Percentages of cells with PLA interactions observed in the nuclei (red; 1 spot/nucleus, dark red; multiple spots/nucleus), and in the cytoplasm (green) normalized to negative controls (mean with SEM, n=4, *p<0.05). (E) UCA1 transcript enrichment in RNA-IP analysis of HT29 cells with or without 2 µM 5-FU for 24h; Input (5% of total), control IgG (defined as 100%) and IP with either BRM or BRG1 antibodies (mean with SEM, n=3).

### Loss of UCA1 results in different SWI/SNF-complexes

To study differences in subunit composition of the SWI/SNF-complexes, we analyzed the interaction of both the ATP-helicases BRM and BRG1 with Bromodomain containing 7 (BRD7) and 9 (BRD9) proteins in our cell model treated with 5-FU. BRD7 is present in BRG1-exclusive PBAF complexes, ^22–24^ and, indeed, our PLA analysis showed no interaction of BRD7 with BRM in ICP cells (Figure 6A). However, such interaction was observed in the dUCA1 cells (Figure 6A). BRD9 is present in embryonic stem cell, non-canonical and progenitor BAF complexes (BRM/BRG1), ^21,29,38,39^ and our PLA results showed an interaction of BRD9 with BRM in the ICP cells, which increased in the dUCA1 cells (Figure 6B). The interaction of BRD7 with BRG1, related to canonical PBAF complexes, was observed in the ICP cells, but was decreased in the dUCA1 cells (Figure 6C). On the contrary, the interaction of BRD9 with BRG1 (BAF complexes) was low in ICP cells, but was high in the dUCA1 cells (Figure 6D). Taken together, our PLA interaction results suggest that UCA1 interaction with BRM- and BRG1-containing complexes altered the subunit composition of the SWI/SNF chromatin remodeling machinery. In particularly, loss of UCA1 shifted the balance towards BRD9-BRG1 complexes previously implicated in embryonic stem cells.

**Figure 6.**
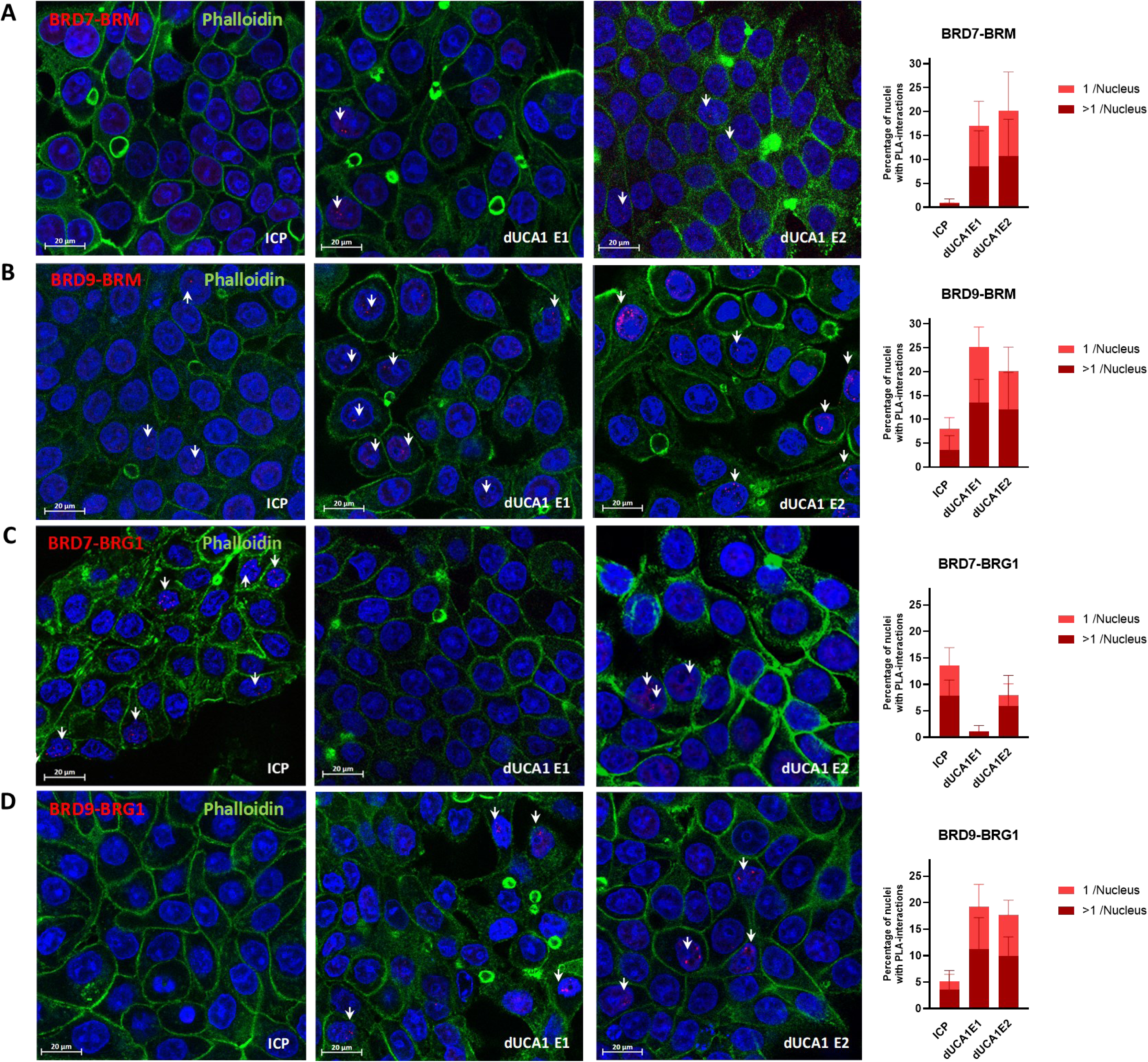
Depletion of UCA1 inverts the balance of different SMARCA4 - SWI/SNF complexes. (A) PLA interactions in control ICP, dUCA1E1 and dUCA1E2 cells with 2 µM 5-FU for 48h (mean with SEM, n=4) for BRD7 and BRM, (B) BRD9 and BRM, (C) BRD7 and BRG1, (D) BRD9 and BRG1 On the left representative confocal microscope images with nuclear staining (Dapi, blue), actin (Phalloduin, green) and dotted interaction signals (PLA, red; indicated by white arrows), and, on the right, PLA quantification depicting percentages of cells with interactions observed in the nuclei (red; 1 spot/nucleus, dark red; multiple spots/nucleus).

## DISCUSSION

We confirmed that lncRNA UCA1 expression is increased in CRC tumor tissues and showed that it is defined to a particular subclass of tumor cells. In the colorectal cancer cell line HT29, UCA1 expression was increased by 5-FU and by oxaliplatin treatment. In addition, we also observed an increased expression upon the DNA damage induced by CRISPR/CAS9 transfection in these cells. CRISPR/CAS9 transfection has recently been described to induce p53-activation in several cancer cell lines ^40^. Interestingly, UCA1 also binds several miRNAs of the p53 pathway^4^ and may play a role in this activation. However, this does affect our analysis, since we used HT29 cells with inducible Cas9 in our studies. Hence, we observed a decreased UCA1 expression after multiple passaging of HT29iCas9 cells without Cas9 induction showing that the increased UCA1 level after transfection was transient.

Our study showed that in those tumor cells that express UCA1, this lncRNA can interfere with the assembly of specific SWI/SNF chromatin remodeling complexes. Firstly, UCA1 depletion in HT29 cells led to an increase of cells with a stem cell phenotype and lower levels of SMARCA2 in cell expressing the stem cell markers CD44, CD166 and CD133. Moreover, after treatment with 5-FU or oxaliplatin, we showed that the increased UCA1 expression is permissive for transcriptionally upregulating of SMARCA2 expression, which compensates for the BRM (*SMARCA2*) protein loss. It is known that BRM may restitute BAF complex functionality in the absence of BRG1, but that it cannot universally compensate for BRG1 function.^41,42^ BRG1 is present in (non-)/canonical BAF complexes and it is the exclusive ATP helicase of PBAF complexes. Previously, inhibition of BRM-BAF complexes by lncBRM was reported to stimulate the formation of BRG1-BAF complexes and stimulated stemness.^43^ Thus, similarly, UCA1-stimulated SMARCA2 transcription may function as a molecular switch between SWI/SNF activating and repressing chromatin binding complexes.

Secondly, we here showed that UCA1 physically interacted with the two ATP-helicases BRM (SMARCA2) and BRG1 (SMARCA4) in the nucleus of the colorectal HT29 cells, suggesting that UCA1 may also physically interfere with the SWI/SNF subunit assembly. Such effects have been described for other lncRNAs, e.g. the action of lncRNA NEAT on the balance of BRD4’s activator and repressor chromatin complexes.^44^ Indeed, our results showed that in the absence of UCA1 the interaction of BRD7 and BRG1 is decreased, and that of BRD9-BRG1 is stimulated. BRD7 is present in BRG1-exclusive PBAF complexes, whereas BRD9 is present in non-canonical and progenitor BAF complexes (BRM/BRG1).^21–24,38^ The exact composition of the BAF complex requires further investigations. SWI/SNF complexes have a dynamic composition and it is not always clear in literature which subunits are present in the different complexes. For example, it is unclear if BRD9 can be part of the canonical BAF complex. We here showed interaction of BRD9 with BRG1 suggesting it is in such complexes. The observed increase in stem cell properties in dUCA1 cells could be explained by loss of PBAF, frequently associated with differentiated cells, and concomitant increase of canonical BRG1-BAF complexes, such as esBAF.

In addition, the BRM-associated complexes that are increased with UCA1 could explain other observations; when HT29iCas9-dUCA1 cells were seeded in high cell concentrations for sphere-culture, the spheres tended to merge. It is reported that BRM may regulate the formation of tight junctions and UCA1 can stabilize E-cadherin in prostate cancer.^4^ Therefore, further analysis of tight-junction factors may reveal a role of UCA1, via BRM, in cell contact regulation.

Recent studies focused on vulnerabilities of cancers presenting mutations of SWI/SNF-encoding genes and the use of proteolysis-targeting chimeras to target the SWI/SNF complexes.^34,35^ It would be interesting to target not only the protein complexes, but also the lncRNAs that may be associated and that may coordinate assembly/stabilization of certain SWI/SNF complexes implicated in cancer aggressiveness.

In conclusion, our study showing that UCA1 interacts with the ATP-helicases from the SWI/SNF chromatin remodeler complexes and influences the presence of different SWI/SNF chromatin remodeler complexes, thereby restraining cells to gain stem cell properties, highlights the importance of lncRNAs vis-à-vis the assembly of protein complexes.

## List of abbreviations

AAVS1: Adeno-Associated Virus Integration Site 1
ARID1A: AT-Rich Interaction Domain 1A protein (BAF250A)
BAF: BRG1-or BRM-Associated Factors
BICeL: “Plateformes Lilloises en Biologie et Santé“ (PLBS) - UAR 2014 – US 41
BRG1: Brahma related gene 1 protein (encoded by SMARCA4)
BRM: Brahma protein (encoded by SMARCA2)
cBAF: canonical BAF
ncBAF: non-canonical BAF
CRCs: colorectal cancers
ELDA: Extreme limited dilution assay
FDR: False discovery rate
5-FU: 5-Fluorouracil
HR: hazard ratio
iCas9: inducible CRISPR associated protein 9
lncRNA: long non-coding RNA
NAT: Tumor-adjacent normal tissue
PBAF: PolyBromo-Associated Factors
PLA: Proximity Ligation assay
RNA-IP: RNA-Immuno-Precipitation
RNAseq: next-generation RNA sequencing
SMARC-A/B/C/D: SWI/SNF related, matrix associated, actin dependent regulator of chromatin, subfamily A /B/C/D (used to refer to gene transcripts)
TCGA: The Cancer Genome Atlas Program

## ACKNOWLEDGEMENTS

We are indebted to all individuals who participated in the patient study from the « Chirurgie Générale et Digestive » and « Chirurgie Digestive et Transplantation » of the Lille university hospital (OrgaRES consortium study) and we acknowledge the Lille ALLIANCE-CANCER Biobank for making the tissue samples available. We acknowledge the technical assistance and expertise of the “Plateformes Lilloises en Biologie et Santé“ (PLBS – UAR 2014 – US 41, Lille, France) with special thanks to Nathalie Jouy (BICeL), Emilie Floquet (BICeL) and Shéhérazade Sebda (Go@L-GFS). We also thank Isabelle Guignon and Helene Touzet for bioinformatics assistance with Cas9 single-guideRNAs (Bilille, PLBS – UAR 2014 – US 41). We acknowledge the technical assistance of undergraduate students Nail Bouzalmad (Université de Bretagne Occidentale, France) and Hélène Marlier (Université de Mons, Belgium).

## Methods

### Key resources table

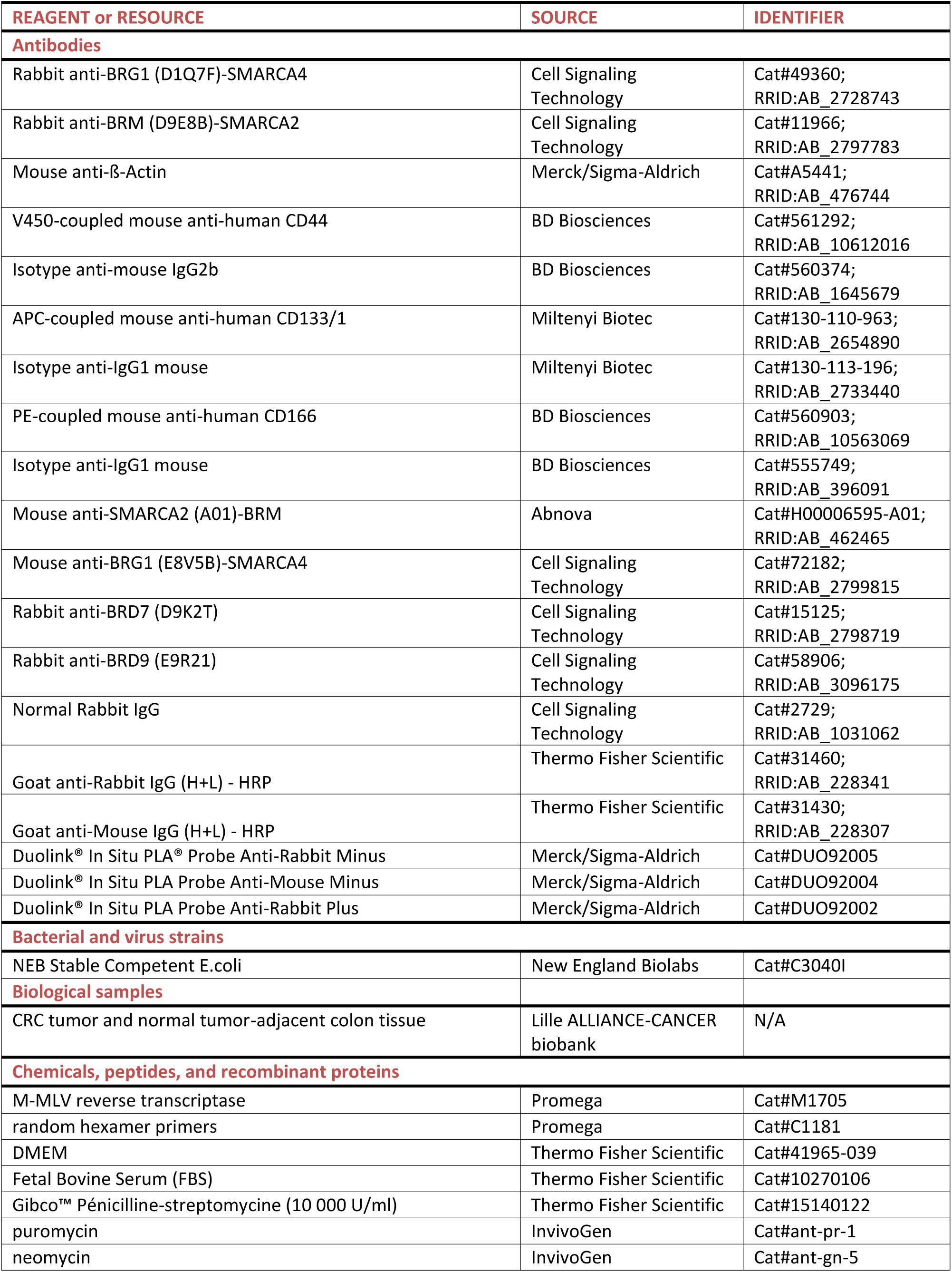
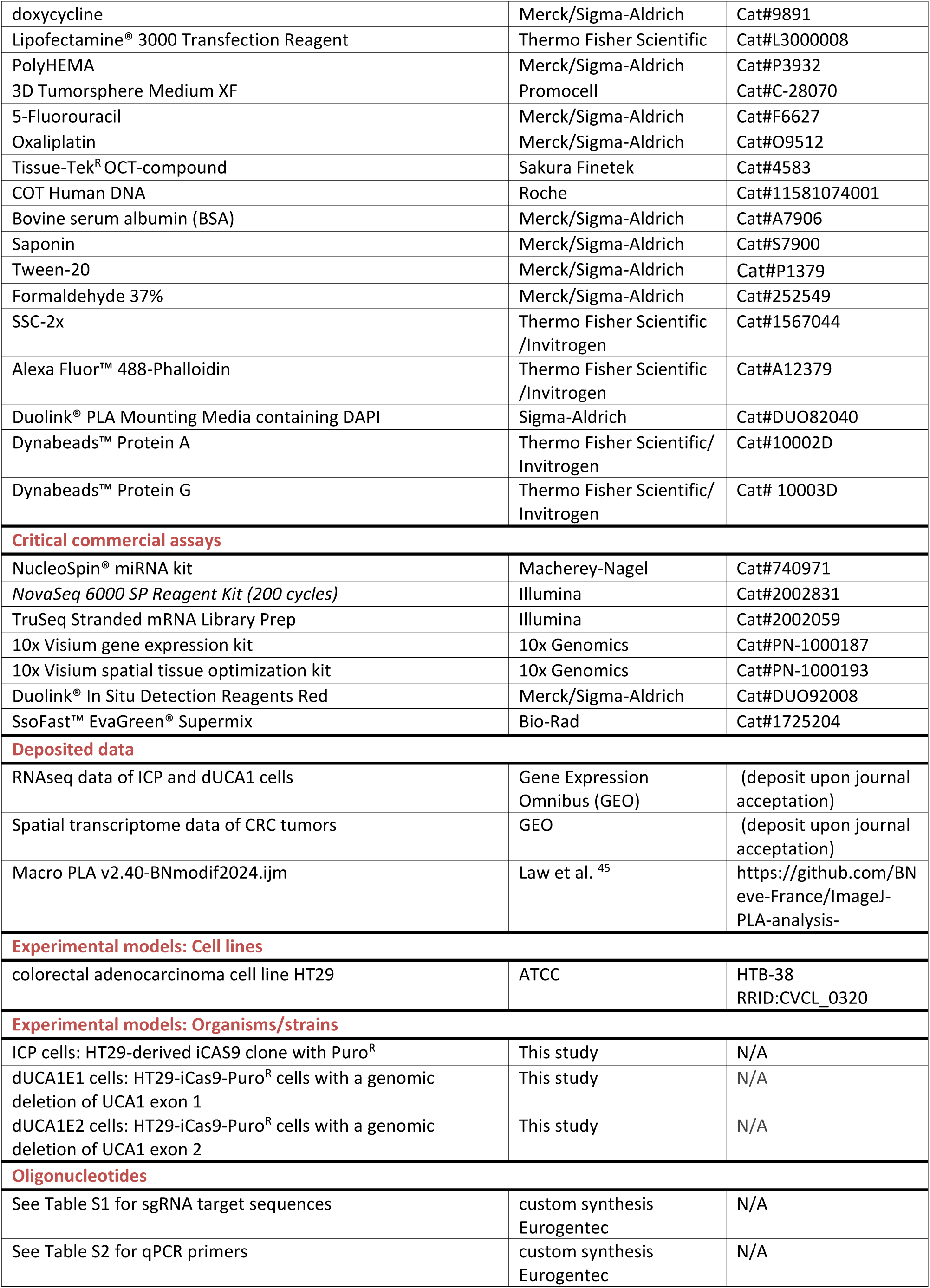
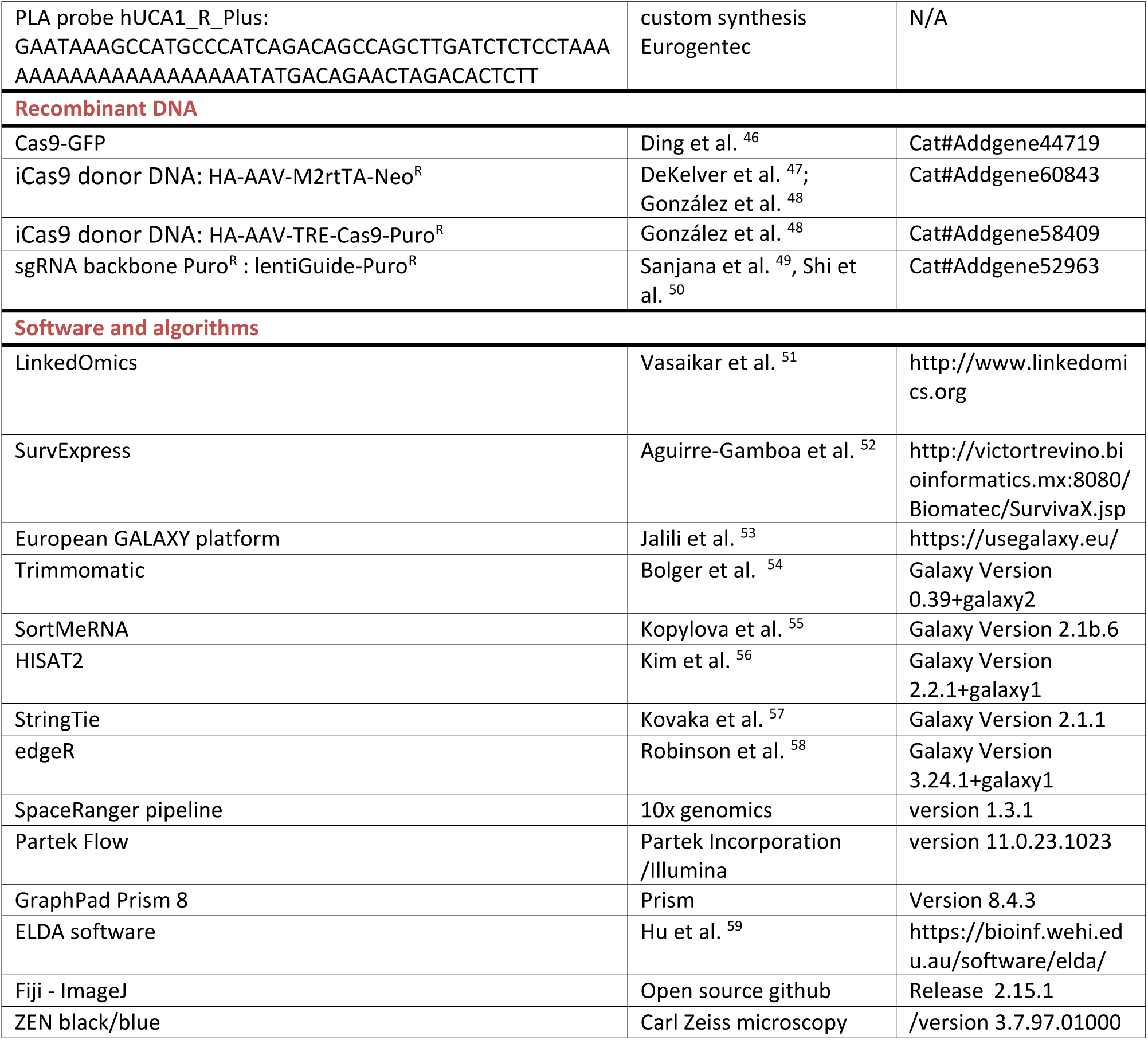

### Resource availability

Further information and requests for resources and reagents should be directed to and will be fulfilled by the lead contact, Bernadette Neve.

### Materials availability

Cell line derivates generated in this study are available from the lead contact upon request.

### Data and code availability

- RNA sequencing and spatial transcriptome data is available on GEO once accepted for journal publication. This paper analyzed existing, publicly available data from the TCGA COAD-READ study (http://www.linkedomics.org) and the Human Colon Cancer Atlas (Single cell expression data of the c295, 62 CRC patients; https://singlecell.broadinstitute.org [29]).
- This paper does not report original codes.
- Any additional information required to reanalyze the data reported in this paper is available from the lead contact upon request.

### Experimental model and study participant details

The OrgaRES consortium study includes CRC tumor and normal tumor-adjacent colon tissues from 13 patients at the university hospital of Lille (CHU Lille, France). Samples were made available by the Lille ALLIANCE-CANCER biobank.

### Method details

#### Study population

The OrgaRES consortium study includes CRC tumor tissues and normal tumor-adjacent colon tissues, from 13 patients at the university hospital of Lille (CHU Lille, France), that were cryopreserved in Tissue-Tek R OCT-compound. All patients gave their informed consent (sex = 9 M; 4 F; mean age at surgery 67 ± 9).

#### Online dataset analysis

Survival data and normalized transcript expressions of the TCGA data sets was assessed via linkedOmics ^51^. The TCGA COAD-READ study included 391 patients histologically diagnosed with colon adenocarcinoma, 152 with rectal adenocarcinoma and 95 diagnosed with mucinous adenocarcinomas. Included data are the “HiSeq RNA” expression (Firehose_RSEM_log2) from Broad Institute of MIT and Harvard (version January 2016). Association of SWI/SNF genes with survival in the TCGA study was assessed via SurvExpress ^52^. Single cell expression data of the Human Colon Cancer Atlas (c295, 62 CRC patients; GSE178341), was assessed using the portal from the Broad Institute of MIT and Harvard (https://singlecell.broadinstitute.org).^60^

#### RNA Expression analysis

RNAs were isolated from 80%-confluent cultured cells and from cryopreserved resected patient samples using the NucleoSpin® miRNA kit. The isolated RNA was reversed transcribed using M-MLV reverse transcriptase and random hexamer primers. Quantitative real-time PCR (qPCR) was performed with SsoFast™ EvaGreen® Supermix on a Bio-Rad CFX96-C1000 Touch Thermal Cycler. To calculate relative expression (2^-ΔΔCt^ method), RPLP0 was used as reference gene and samples were normalized vs control. Primer sequences are listed in Table S1.

Bulk RNAseq and alternative transcript analysis was performed by RNA sequencing of two replicated experiments using the NovaSeq 6000 System (Plateforme de Genotypage et Séquençage, Institut du Cerveau, Paris, France). Data was analyzed using the European GALAXY platform:^53^ Raw paired-end sequencing reads were cleaned using Trimmomatic^54^ and ribosomal RNA reads were filtered out by SortMeRNA.^55^ Cleaned reads were aligned to the human Gencode reference (GRCh38.p13, v36) using HISAT2.^56^ Transcriptome assembly and quantification was applied by StringTie. ^57^ Normalization and differential expression analysis were performed using edgeR. ^58,61^

#### Spatial transcriptome expression

Tumor tissues from CRC patients (OrgaRES consortium study, France) were cryopreserved by snap freeze in Tissue-Tek^R^ OCT-compound and subjected to the 10x Visium gene expression analysis according to the manufacturer guidelines. Prior to the analysis a “spatial tissue optimization” was conducted, which determined the optimal tissue permeabilization time of 30 min. A Hematoxylin-Eosin staining was done after the tissue section was deposited onto the Visium slide capture area, images were obtained by Axioscan (Zeiss) and tumor tissue annotated by a pathologist. After reversed transcription on slide, a library was construction at the Go@L-GFS platform (PBLS - UAR2014 - US41, Lille France). The library was sequenced on a SP flowcell of the NovaSeq 6000 system (Illumina; Table S2). Using data of the SpaceRanger pipeline, gene and differential expression analysis were performed with Partek Flow (version 11.0.23.1023). In short, regularize negative binomial regression was performed for normalization and variance stabilization (SCTransform) and sample batch variation corrected by Seurat 3 integration. Thereafter, principal component and graph-based clusters were analyzed.

#### Derivation of UCA1-depleted HT29 cells using CRISPR/Cas9

The HT29 cell-line was cultured in DMEM supplemented with 10% (v/v) FBS, and penicillin - streptomycin (100 U/mL-100 μg/mL) at 37°C in humidified air containing 5% CO_2_. To obtain cells with inducible Cas9 expression (HT29-iCas9), HT29 cells were lipo-transfected with single-guide (sg)RNA AAVS1 (Table S3), Cas9-GFP and iCas9 donor DNA plasmids from González et al. ^48^ The genomic DNA from the antibiotic resistant HT29-iCAS9 clones was analyzed by Sanger sequencing and the ones without AAVS1 wild type alleles were selected. These cells were cultured in DMEM supplemented with 10% FBS (v/v), 0.5 µg/mL puromycin and 250 µg/mL neomycin. Next, these HT29-iCAS9 cells were incubated with 2 µg/ml doxycycline for 2h before lipofectamine3000 transfection with 2 µg paired sgRNA constructs (Table S3). After 48h, cellular clones were isolated based on their GFP-expression by flow cytometry at the Lille cytometry platform (BD FACS Aria cell sorter, BICeL; “Plateformes Lilloises en Biologie et Santé“ (PLBS) - UAR 2014 – US 41), obtaining HT29-iCas9-transfected (ICP), dUCA1E1 and dUCAE2 cells.

#### Sphere formation analysis

Sphere formation was assessed by cell cultures on non-adherent 96-well plates using either DMEM/F12 without serum and in the presence of growth factors or with 3D-Tumorsphere Medium XF. To determine the presence of cells with stem-cell properties, sphere formation was analyzed using both extreme limited dilution assays (ELDA) as previously described ^59^ and an individual cell culturing assays. For the latter, cells were seeded at 3 cells/well onto a 96-well plate by flow cytometry. With the microscopic system cell discoverer 7 (Zeiss, Marly le Roi, France; BICeL plateform) images of spheres were acquired after 11 days of culture, and size analysis performed with ImageJ software.

#### FACS-based analysis of cells expressing stem cell markers

Suspended HT29 cells were incubated with 1.25 µg fluorophore-coupled antibodies per 10^6^ cells in DMEM culture medium supplemented with 10% (v/v) FBS and 0.5 mM EDTA for 30 min. Used antibodies were V450-coupled mouse anti-human CD44, APC-coupled mouse anti-human CD133/1, PE-coupled mouse anti-human CD166 and their isotype-controls. Thereafter, cells were sorted into a pool of cells expressing no markers (−), one of three (+), two of three (++), or all markers (+++) by flow cytometry using the BD FACSAria cell sorter at the Lille cytometry platform (BICeL). The sorted cells were either recovered in Tumorsphere Medium XF medium or in RNA-lysis buffer for transcript expression analysis.

#### Protein expression

Proteins were recovered during the NucleoSpin® miRNA procedure according to the manufactory’s guidelines or from cell lysates prepared with RIPA buffer. The samples were quantified and submitted to SDS-PAGE using NuPAGE™ Novex™ 4-12% Bis-Tris Protein Gels. Western blot immuno-labelling was performed with a rabbit monoclonal antibody specific for BRG1 (ab110641, 1:10000), BRM (Cell signaling#1966, 1:10000) or ß-Actin mouse antibody (A5441, 1:5000) accordingly to the manufactory’s protocol. Secondary HRP-labeled antibodies (Invitrogen, diluted 1:5000) and SuperSignal™ West Pico PLUS chemiluminescent substrates were used to visualize the proteins with the ImageQuant LAS 4000 camera (Fujifilm Corporation). Quantifications were analyzed with the ImageJ program (https://imagej.net/Fiji).

#### Proximity Ligation assay

Cells were cultured in Ibidi μ-Slides VI 0.4 or 8-chamber slides (Cliniscience, Nanterre, France), fixed with 4% (v/v) formaldehyde and permeabilized with 0.1% (w/v) saponin.

To study lncRNA-protein interactions, we used the Duolink® Proximity Ligation assays (PLA) as previously described ^62^. First, we blocked background signals with 20 μg/mL sheared Cot-DNA in Duolink® blocking solution. Secondly, the slides were incubated with 100nM denatured hUCA1_R_Plus probe in Duolink® blocking solution overnight at 4°C. After three PBS washes, an additional blocking was done with PBS containing 0.1% (v/v) Tween-20, 20 μg/mL sheared Cot-DNA and 1% (w/v) BSA at room temperature for 1h. Slides were washed respectively with PBS, SCC-2x containing 0.1% (v/v) Tween-20, and with PBS. Thereafter, the manufacturer instructions were followed using the primary rabbit antibody BRG1 (1:250), BRM (1:250) or normal Rabbit IgG control (1:250) and the secondary antibody Duolink® In Situ PLA® Probe Anti-Rabbit Minus.

To study protein-protein interactions, we followed the PLA protocol provided by the manufacturer using primary mouse antibodies BRG1 (1:350) and BRM (1:250), rabbit antibodies BRD7 (1:250) and BRD9 (1:250) in combination with the secondary antibodies Duolink® In Situ PLA® Probe Anti-Rabbit Plus and Duolink® In Situ PLA Probe Anti-Mouse Minus. Control incubations with one primary antibody were performed in parallel.

The PLA signal was revealed by the Duolink in situ detection Red kit. After the washes with Duolink® wash buffer B, the slides were incubated with 1:1000 diluted Alexa Fluor™ 488-Phalloidin in wash buffer B for 20 min, washed with buffer B and mounted with Duolink® PLA Mounting Media containing DAPI. Images were acquired using a confocal microscope (LSM 710, Zeiss) and the PLA-interaction signals were counted as discrete fluorescent spots (ƛ 594 nm) in maximal intensity projections of the whole nucleus using the modified ImageJ macro “PLA v2.40” ^45^ (PLA v2.40-BNmodif2024.ijm). At least 100 nuclei per experimental condition were analyzed and statistics was performed using Prism 8 (n=4).

#### RNA-Immuno-Precipitation assay

RNA-Immuno-Precipitation (RNA-IP) was performed as previously described ^63^. Briefly, 80% confluent HT29 cells were crosslinked with a 1% (v/v) formaldehyde solution in PBS for 15 min, and this was quenched with 125 mM glycine. Cells were washed with PBS, collected by centrifugation and there after the nuclear fraction was isolated and the chromatin was sheared by sonification to 200-500bp fragments using Bioruptor (Diagenode). Overnight immuno-precipitation was performed with either 1:100 diluted rabbit monoclonal antibody BRG1, rabbit monoclonal antibody BRM or normal Rabbit IgG control. To precipitate the immune complexes a 30 µL mixture of Protein A/G Dynabeads (1:3) was used. After several washes the bound fraction was eluted and incubated 2h at 80°C for decrosslinking. Proteins were degraded by incubation with 500 µg/mL proteinase K for 1h at 55°C, where after RNA was isolated using NucleoSpin® RNA kit in the presence of carrier RNA.

### Quantification and statistical analysis

For RNA transcript and protein expression analysis GraphPad Prism 8 was used to apply paired t-tests on ratios to compare UCA1 expression in NAT vs Tumor tissues, and to apply One-way ANOVA followed by multiple comparison tests for cell line experiments. Figures represent mean with SD, unless stated differently and the number of biological replicates and statistically significant results (*p*-values) are detailed in the figure legends. For bulk RNA-seq analysis, a pairwise comparison test was performed using EdgeR at the European GALAXY platform.^53,58,61^ Partek Flow was used to assess gene correlation and gene set enrichment in the spatial transcriptome data. In short, regularize negative binomial regression was performed for normalization and variance stabilization (SCTransform) and sample batch variation corrected by Seurat 3 integration. Thereafter, principal component analysis and graph-based clusters were analyzed. The ELDA software provided the pairwise tests for differences in stem cell frequencies (χ2).^59^ With GraphPad Prism 8 survival curves and statistical analysis of averaged expression data from TCGA were performed (Mantel-Cox log-rank hazard ratio (HR) and P-value (*p*)).

## DECLARATIONS

### Ethical Approval

The OrgaRES consortium study includes tissues samples from patients at the university hospital of Lille (CHU Lille, France) obtained with the approval of the institutional review board and the Lille ALLIANCE-CANCER biobank. All patients gave their informed consent and signed a non-opposition statement to research use of a biological sample.

### Authors’ contributions

BN designed and conducted the research, analyzed and interpreted the data, and wrote the article. EHB conducted the research, analyzed and interpreted the data and edited the article. BD conducted the research on human tissue samples. MS conducted the research, analyzed and interpreted the data. MF organized the preparation and realization of the Visium Library sequencing and JPM analyzed the spatial gene expression (SpaceRanger/Loupe). EL and the OrgaRES consortium provided histology expertise and human tissue samples. NJ provided expertise and analysis of TCGA data, and edited the article. AV discussed results and experimental setup, reviewed and edited the article. IVS provided expertise and revised the article. All authors read and approved the final manuscript.

