## Supplementary material for "Long non-coding RNA UCA1 affects chromatin remodeling via SMARCA2-containing SWI/SNF complex in human colorectal cancer": BNeve et al 2024_UCA1-BAF_Supplement.pdf

### Legends to supplementary figures

##### Figure S1. RNAseq of ICP and dUCA1 cells.

(A) Schematic representation of detected UCA1 variants (based on Ensembl; https://www.ensembl.org). Depicted by red arrows are the positions of the sgRNAs targeting exons of UCA1. Alternative transcripts that can be detected by the qPCR primers are indicated with blue dots. HT29-derived cellular clones (control ICP, dUCA1E1 and dUCA1E2 cells) were treated for 72h with or without 2 µM 5-Fluorouracil (5-FU) and analyzed by RNAseq.

(B) Heatmap of alternative UCA1 transcript expressions

(C) Heatmap depicting log Fold-Changes (FC) of UCA1 target genes from literature compared to control ICP treated with 5-FU

##### Figure S2. Correlation of SWI/SNF subunits with UCA1.

(A) The TCGA data of colon adenocarcinoma (COAD),

(B) pancreatic adenocarcinoma (PAAD),

(C) esophageal carcinoma (ESCA),

(D) Stomach adenocarcinoma (STAD) and

(E) Stomach and esophageal carcinoma (STES)t.

Data was analyzed with linkedOmics. Correlation with UCA1 expression was assessed for SMARCA2 and SMARCA4. Pearson correlation coefficients (r) and p-values (*p*) were calculated using GraphPad Prism 8.

##### Figure S3. RNA transcript expression after oxaliplatin treatment in dUCA1 cells.

(A) UCA1 transcript expression of control ICP, dUCA1E1 and dUCA1E2 cells treated for 72h with 2 µM oxaliplatin (OXA).

(B) SMARCA2 transcript expression

(C) SMARCA4 transcript expression

(n=3, *$p < 0.05$).

### Supplementary tables

#### Table S1. Sequence of qPCR primers

| qPCR primers | |
|---|---|
| hRPLP0ex7_F | GCAATGTTGCCAGTGTCTG |
| hRPLP0ex7-8_R | GCCTTGACCTTTTCAGCAA |
| hUCA1ex2_F | GCCCCTTGGACCATCACAG |
| hUCA1ex3_R | GTTTGAGGGGTCAGACTTTTGA |
| hSMARCA2ex7_F | CAGAAGATTGAGCAGGAGAGGAAAC |

| | |
|---|---|
| hSMARCA2ex9_R | ATTGGCTACATACTCATCGGTCTGC |
| hSMARCA4ex4_F | GAGTCCATGCATGAGAAGGG |
| hSMARCA4ex5_R | GAACTGGACTAGAGGCATGC |

#### Table S2. Visium - Space ranger summary

| Patient | Visium Tissue Spots (of 5000) | Sequencing depth (Mean reads/spot) | Median genes (genes/spot) |
|---|---|---|---|
| 1 | 2771 | 128 962 | 4 243 |
| 2 | 2230 | 161 888 | 3 804 |
| 3 | 947 | 200 636 | 7 379 |
| 4 | 2194 | 168 632 | 6 064 |
| 5 | 2315 | 142 043 | 2 870 |
| 6 | 2465 | 61 091 | 3 224 |
| 7 | 2670 | 92 221 | 2 463 |
| 8 | 1809 | 215 503 | 4 098 |
| 9 | 2715 | 60 288 | 3 709 |
| 10 | 2224 | 130 655 | 3 302 |
| 11 | 1691 | 247 054 | 1 968 |
| 12 | 2735 | 71 100 | 2 336 |

#### Table S3. Target sequences of sgRNA

| sgRNA | Target-specific sequence (5'-3') |
|---|---|
| AAVS1[47] | GGGGCCACTAGGGACAGGAT |
| UCA1 Exon 1 upstream | ATCTCCCTCCGTCATCTAAA |
| UCA1 Exon 1 downstream | AGACGTTTGAGTTTTATATC |
| UCA1 Exon 2 upstream | GGTTTTAAAGGAGTGCCTAG |
| UCA1 Exon 2 downstream | GCGAAAGAGGCCTTGAATTG |

### Figure S1

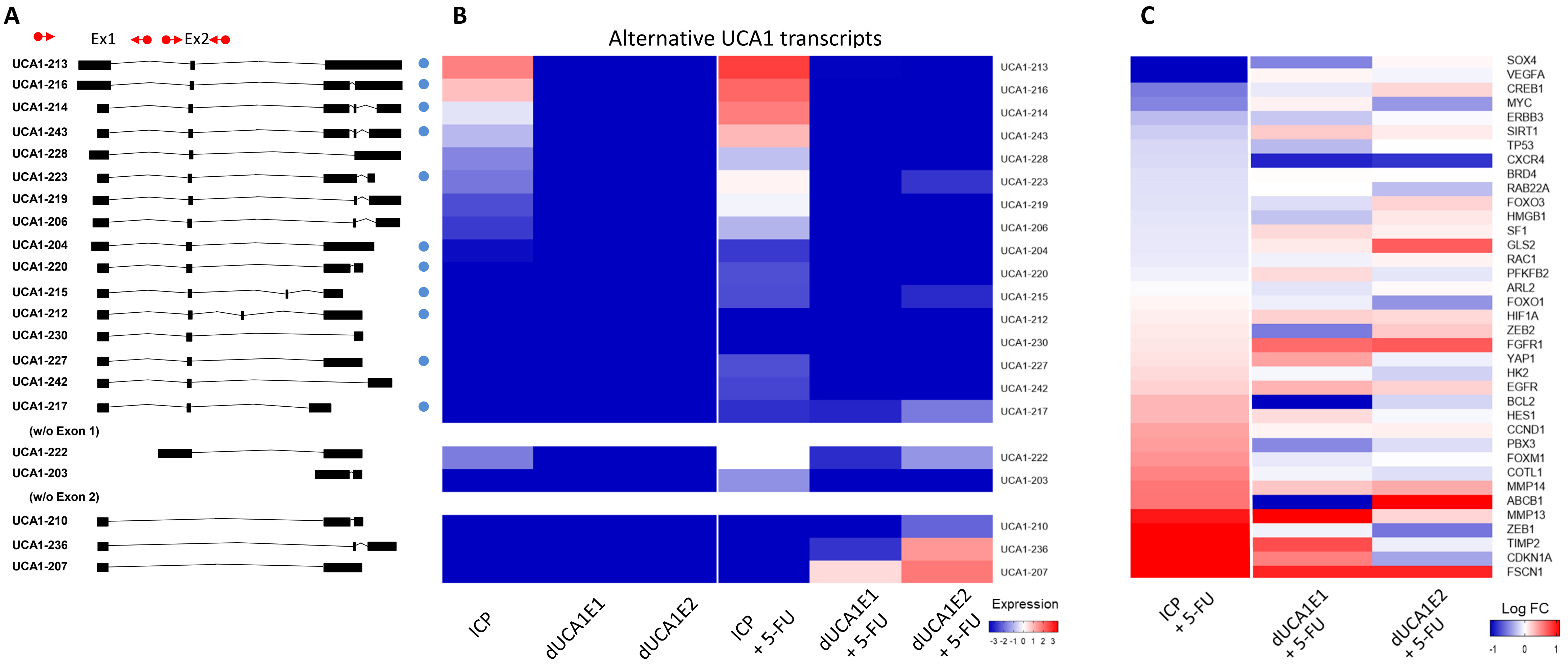

A
Ex1
Ex2
UCA1-213
UCA1-216
UCA1-214
UCA1-243
UCA1-228
UCA1-223
UCA1-219
UCA1-206
UCA1-204
UCA1-220
UCA1-215
UCA1-212
UCA1-230
UCA1-227
UCA1-242
UCA1-217
(w/o Exon 1)
UCA1-222
UCA1-203
(w/o Exon 2)
UCA1-210
UCA1-236
UCA1-207
B
Alternative UCA1 transcripts
ICP
dUCA1E1
dUCA1E2
ICP + 5-FU
dUCA1E1 + 5-FU
dUCA1E2 + 5-FU
Expression
-3 -2 -1 0 1 2 3
C
SOX4
VEGFA
CREB1
MYC
ERBB3
SIRT1
TP53
CXCR4
BRD4
RAB22A
FOXO3
HMGB1
SF1
GLS2
RAC1
PFKFB2
ARL2
FOXO1
HIF1A
ZEB2
FGFR1
YAP1
HK2
EGFR
BCL2
HES1
CCND1
PBX3
FOXM1
COTL1
MMP14
ABCB1
MMP13
ZEB1
TIMP2
CDKN1A
FSCN1
ICP + 5-FU
dUCA1E1 + 5-FU
dUCA1E2 + 5-FU
Log FC
-1
0
1

### Figure S2

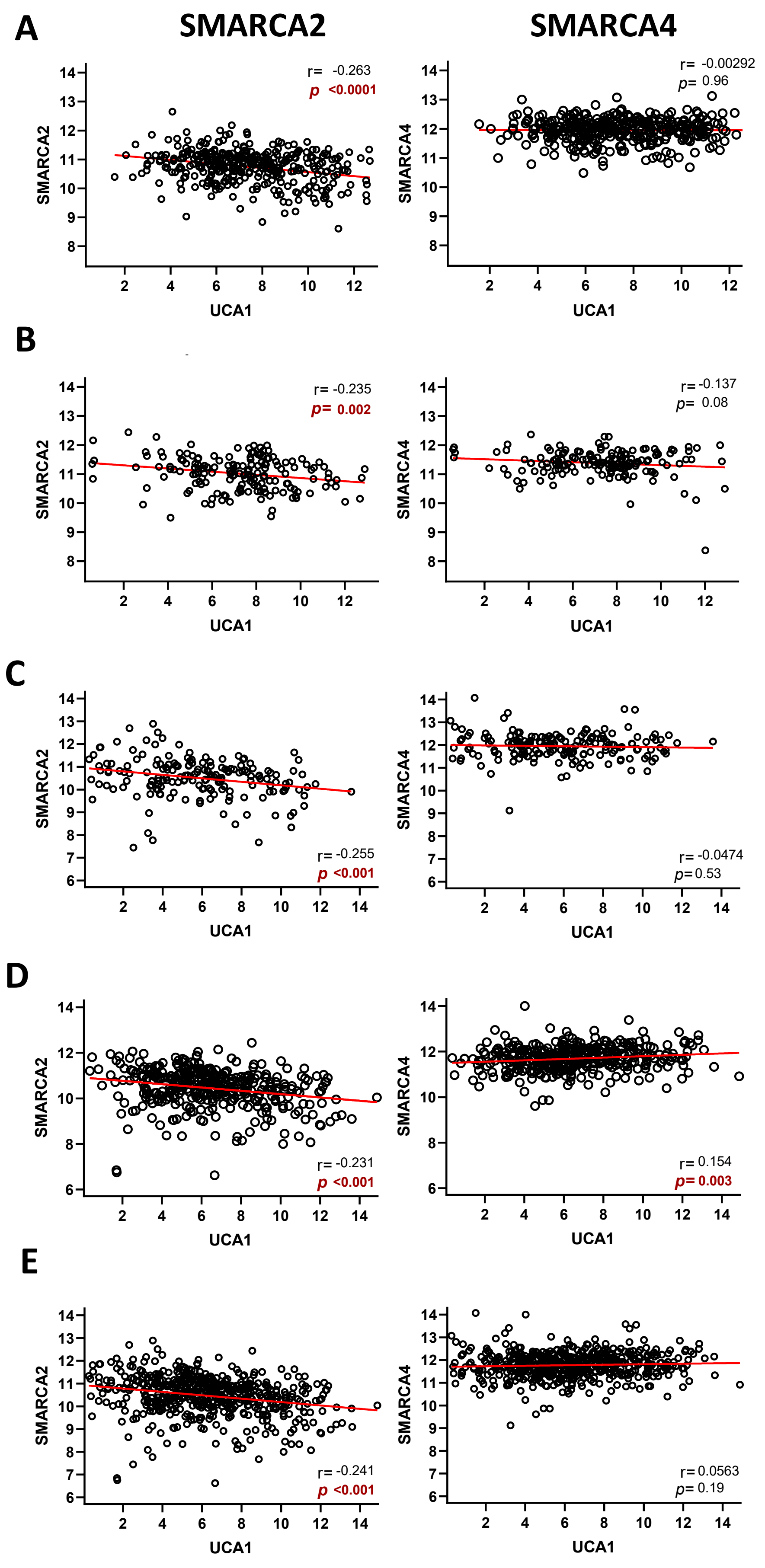

A
SMARCA2
SMARCA4
r= -0.263
p <0.0001
r= -0.00292
p= 0.96
B
r= -0.235
p= 0.002
r= -0.137
p= 0.08
C
r= -0.255
p <0.001
r= -0.0474
p= 0.53
D
r= -0.231
p <0.001
r= 0.154
p= 0.003
E
r= -0.241
p <0.001
r= 0.0563
p= 0.19
SMARCA2
SMARCA4
UCA1

### Figure S3

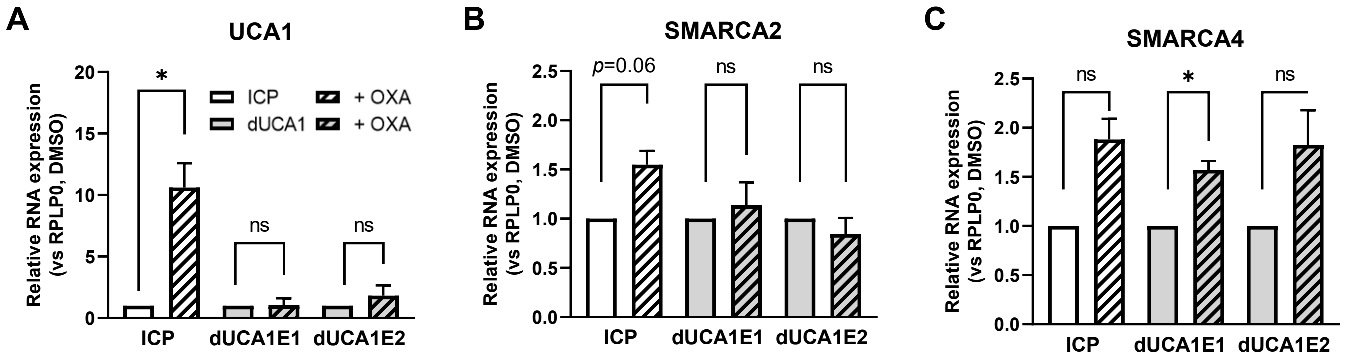
A
UCA1
Relative RNA expression (vs RPLP0, DMSO)
20
15
10
5
0
*
ns
ns
ICP
dUCA1
+ OXA
+ OXA
ICP
dUCA1E1
dUCA1E2
B
SMARCA2
Relative RNA expression (vs RPLP0, DMSO)
2.5
2.0
1.5
1.0
0.5
0.0
p=0.06
ns
ns
ICP
dUCA1E1
dUCA1E2
C
SMARCA4
Relative RNA expression (vs RPLP0, DMSO)
2.5
2.0
1.5
1.0
0.5
0.0
ns
*
ns
ICP
dUCA1E1
dUCA1E2
